# Non-Canonical Functions of a Mutant TSC2 Protein in Mitotic Division

**DOI:** 10.1101/2022.12.07.519401

**Authors:** Mary-Bronwen L. Chalkley, Rachel B. Mersfelder, Maria Sundberg, Laura Armstrong, Mustafa Sahin, Rebecca A. Ihrie, Kevin C. Ess

## Abstract

Tuberous Sclerosis Complex (TSC) is a debilitating neurodevelopmental disorder characterized by a variety of clinical manifestations including epilepsy, autism, and intellectual disability. TSC is caused by mutations in the *TSC1* or *TSC2* genes, which encode the hamartin/tuberin proteins respectively. These proteins function as a heterodimer that negatively regulates mechanistic Target of Rapamycin Complex 1 (mTORC1). TSC research has focused on the effects of mTORC1, a critical signaling hub, on regulation of diverse cell processes including metabolism, cell growth, translation, and neurogenesis. However, non-canonical functions of TSC2 are not well studied, and the potential disease-relevant biological mechanisms are not well understood. We observed aberrant multipolar mitotic division, a novel phenotype, in *TSC2* mutant iPSCs. The multipolar phenotype is not meaningfully affected by treatment with mTORC1 inhibition, suggesting that multipolar division is an mTORC1-independent phenotype. We further observed dominant negative activity of the mutant form of TSC2 in producing the multipolar division phenotype. These data expand the knowledge of TSC2 function and pathophysiology which will be highly relevant to future treatments for patients with TSC.

**Highlights:**

- Novel multipolar division in patient-derived iPSCs with mutant form of tuberin, TSC2 encoded protein
- Mutant tuberin may act in a dominant negative, mTORC1-independent manner

**eTOC:** Tuberous sclerosis complex (TSC) is a disorder caused by mutations in *TSC1* or *TSC2* genes leading to mTORC1 hyperactivity. Chalkley and colleagues found that a mutant microdeletion allele of TSC2 causes multipolar division in human induced pluripotent stem cells. Chalkley and colleagues also found that the multipolar division from mutant TSC2 may have a dominant negative mechanism and be mTOR1-independent.

## INTRODUCTION

Tuberous Sclerosis Complex is a genetic disorder with a broad spectrum of phenotypes, affecting virtually every organ system (Ess, 2010). This disorder occurs in approximately 1 in 6,000 people and is marked by mutations in *TSC1* or *TSC2* which code for hamartin and tuberin, respectively (Identification and characterization of the tuberous sclerosis gene on chromosome 16, 1993; Crino et al., 2006; Sancak et al., 2005; van Slegtenhorst et al., 1997). Loss of function mutations of either *TSC1* or *TSC2* are sufficient to cause pathogenesis, though mutations in *TSC2* have been associated with more severe phenotypes (Sancak *et al*., 2005). Though an autosomal dominant inherence pattern is possible, *de novo* mutations are much more common, and are seen in the majority of patients with TSC (Dabora et al., 2001). Current models of TSC pathogenesis suggest a possible two-hit hypothesis wherein a loss-of function mutation is required in both copies of either *TSC1* or *TSC2* in order to produce disease (Parry et al., 2001). However, not all hamartomas found in patients with TSC reliably demonstrate loss of heterozygosity, suggesting that haploinsufficiency, post-translational inactivation of remaining protein, or dominant negative activities of mutant TSC1 (hamartin) or TSC2 (tuberin) may also underlie some disease phenotypes (Henske et al., 1996; Jozwiak et al., 2008; Neuman and Henske, 2011).

Hamartin and tuberin are canonical negative regulators of the essential mammalian/mechanistic Target of Rapamycin (mTOR) pathway (Dibble et al., 2012; Ess, 2010). Hamartin and tuberin are constitutive inhibitors of RHEB GTPase (Ras homolog enriched in the brain), an upstream regulator of mTOR complex 1 (mTORC1) (Costa-Mattioli and Monteggia, 2013; Inoki et al., 2003; Tee et al., 2003). The principal identified effect of *TSC1* and *TSC2* loss of function mutations is thus unchecked overactivity of mTORC1 (Costa-Mattioli and Monteggia, 2013). mTORC1 promotes protein synthesis, cell growth, and proliferation through phosphorylation of downstream targets, classically including the activation of S6 kinase (S6K) and the inhibition of eukaryotic initiation factor 4E-binding proteins (4E-BP)(Brunn et al., 1997; Burnett et al., 1998; Costa-Mattioli and Monteggia, 2013; Gingras et al., 1999; Hara et al., 1997; Liu and Sabatini, 2020; Ruvinsky et al., 2005).

A large proportion of patients with TSC present with epilepsy and are found to have extensive cortical hamartomas (also known as cortical tubers) of the brain in which lamination is disordered and dysmorphic neurons are present (Ess, 2010). A diagnostic characteristic of cortical tubers is the presence of “giant cells”, so named due to their large size. Giant cells are thought to have abnormal differentiation and can exhibit both astrocytic and neuronal features. Though the cell of origin of giant cells remains controversial, it has been repeatedly observed that giant cells can be multinucleated, a phenotype that can arise from multipolar division (Crino, 2013; Feliciano, 2020; Mizuguchi, 2007; Zarei et al., 2002).

During cell culture experiments with induced pluripotent stem cells (iPSC) derived from a patient with TSC, we observed a high incidence of multipolar dividing cells. This suggests a possible mechanism that may lead to the multinucleated giant cells found in cortical tubers. In this report, we present evidence that a TSC2 loss of function mutant, containing a 6 amino acid deletion within the C-terminus, exhibits a dominant negative function leading to multipolar spindle formation. Abnormal cell division may then produce cells with an atypical distribution of genetic material. This finding highlights a possible contributing factor to the disruptions of normal neuronal migration and cortical development seen in patients with TSC.

## RESULTS

### Patient-derived *TSC2* mutant iPSCs display multipolar division

We have been using an allelic series of isogenic iPSCs derived from a patient with a C terminal 6 amino acid in-frame microdeletion (c.5238_5255del, p. His1746_Arg1751del) in exon 41 of *TSC2* (Sundberg et al., 2018). An accompanying set of isogenic cells was generated from the heterozygous patient-derived iPSC (TSC2 +/LOF) in which the second, wild type allele was also mutated (TSC2 LOF/LOF). Additionally, in a third set of cells the heterozygous mutant allele was corrected to wild type (WT) as previously described (Sundberg *et al*., 2018). A mutant TSC2 protein is produced from the allele harboring the 6 aa in frame microdeletion in the TSC2 +/LOF and TSC2 LOF/LOF cells, although the expression level is decreased relative to TSC2 levels seen in other wild-type iPSCs and the corrected patient-matched line (Figure 1A, B).

**Figure 1:**
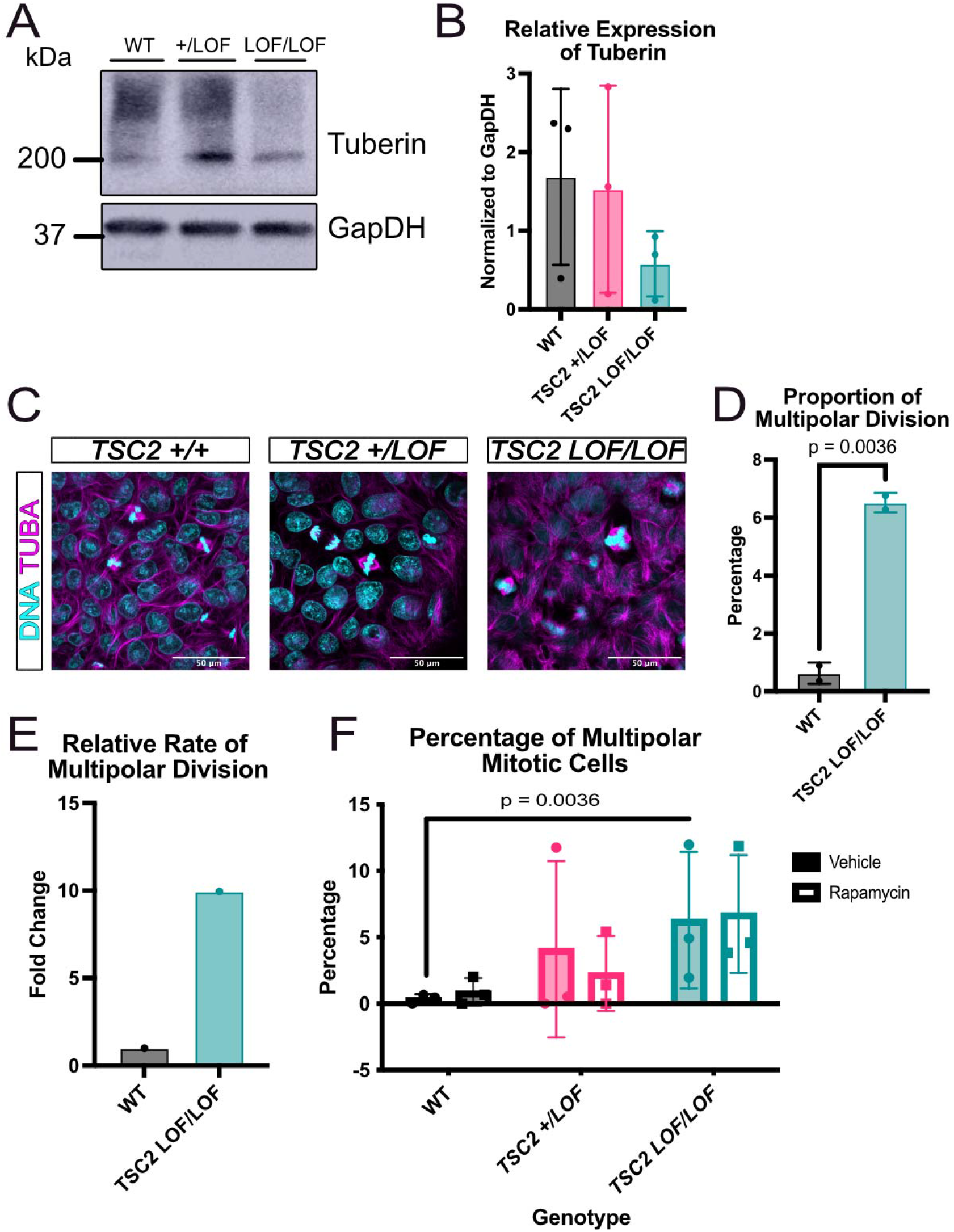
Patient-derived TSC2 loss of function mutant induced pluripotent stem cell cultures have multipolar mitotic cells. (A) Representative immunoblot showing protein expression of tuberin in patient iPSCs. (B) Quantification of relative expression of tuberin (across 3 technical replicates using these lines). (C) Representative immunofluorescence images of induced pluripotent stem cells using TUBA (magenta) to identify mitotic cells and Hoechst (teal) to identify DNA. Scale bars = 50 µm. (D) Average proportion of multipolar dividing nuclei by genotype. N = 3 independent replicates per genotype, p = 0.0036 [student’s t test]. Error bars = SD. (E) Quantification showing relative rate of multipolar division. (F) Quantification showing average percentage of mitotic cells that have the multipolar division per replicate. N = 3 independent replicates per genotype, p = 0.0036, [paired student’s t test]. Error bars = SD.

During standard cell culture maintenance, we observed that patient-derived TSC2 +/LOF iPSC cells exhibited a higher incidence of multipolar division, with cells including three or four mitotic poles as opposed to the expected two poles during mitosis. This phenotype was detected in both the original patient-derived line as well as the homozygous LOF line (Figure 1C). To determine the rate of multipolar division in the mutant patient iPSC line, we counted the total nuclei present and then scored the total number of nuclei undergoing division within this population. The average rate of division was not significantly different in the wild type compared to mutant iPSCs (TSC2 +/LOF and TSC2 LOF/LOF) (data not shown). Nuclei undergoing division and exhibiting more than two spindle poles were considered to be dividing multipolar, and the total number counted. A significant increase in multipolar nuclei as a percentage of dividing nuclei was observed in TSC2 LOF/LOF iPSCs compared to matched wild type iPSCs (p=0.0036) (Figure 1D, E). TSC2 +/LOF heterozygous mutant iPSCs showed an increase in multipolar nuclei compared to wildtype (Figure 1F). To determine if the multipolar phenotype was due to increased mTORC1 activity, wild type and mutant iPSCs were treated with rapamycin, a potent mTORC1 inhibitor. No differences in the percentage of mitotic cells were observed in vehicle treated vs rapamycin treated iPSCs. Interestingly, TSC2 mutant iPSCs (TSC2 +/LOF and TSC2 LOF/LOF) treated with 0.4 nM rapamycin for 24 hours continued to show increased number of multipolar nuclei (Figure 1F).

### Multipolar division phenotype is not observed in TSC2 knock out iPSCs

We sought to replicate these observations in another iPSC line. To test the impact of the presence or absence of tuberin, we used a CRISPR/Cas9 engineered TSC2 knock out iPSC line that did not originate from a patient and produces no tuberin protein ((Armstrong et al., 2017) (Figure 2 A, B). When repeating the approach shown in Figure 1 with this line, the average rate of division and average percentage of dividing nuclei were not significantly different between the isogenic wild type and knock out genotypes (data not shown). Moreover, few to no nuclei were multipolar in the TSC2 homozygous complete knock out iPSCs (Figure 2C, D).

**Figure 2:**
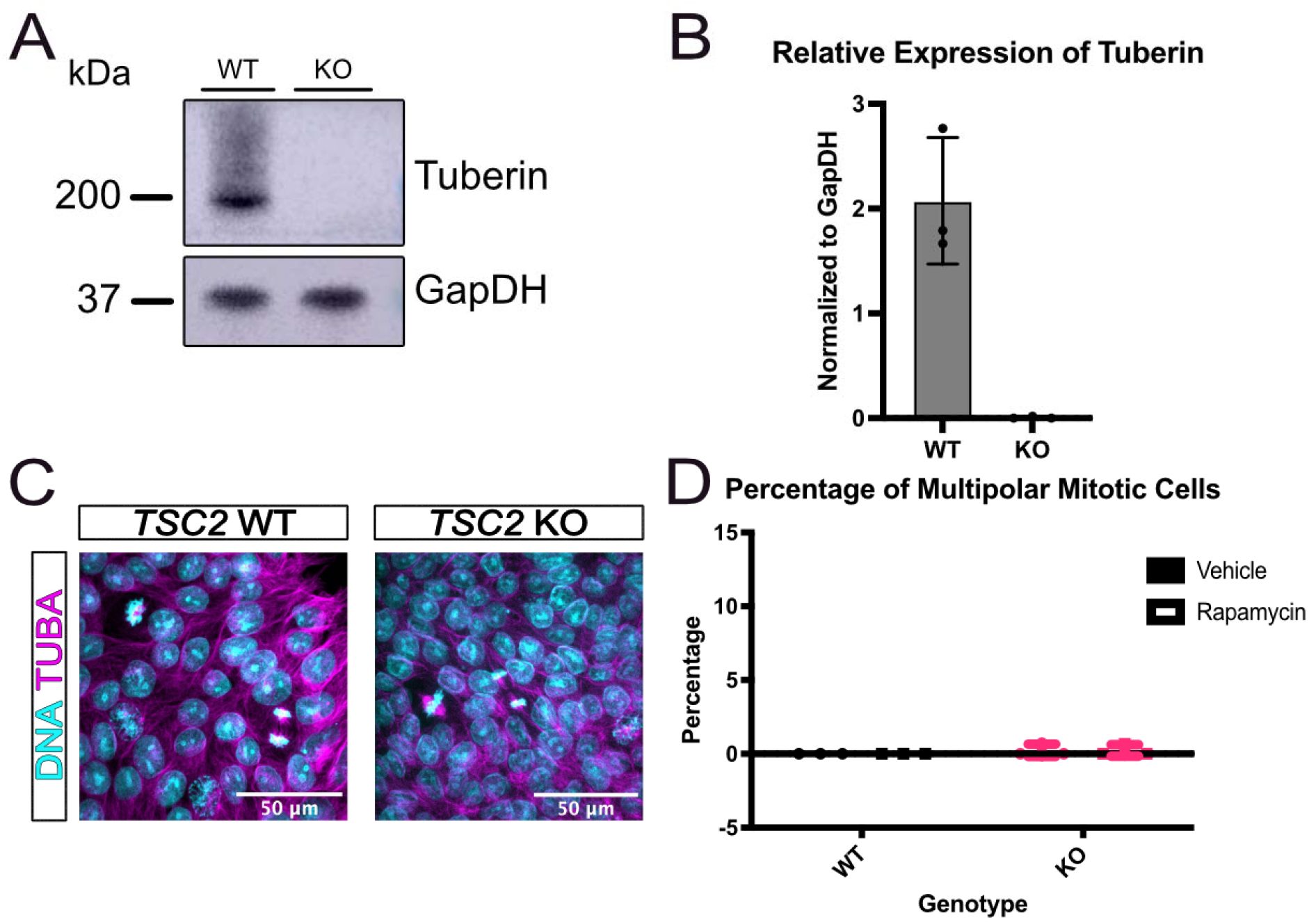
TSC2 knock out induced pluripotent stem cell cultures have no multipolar mitotic cells. (A) Representative immunoblot showing tuberin expression in TSC2 knock out iPSCs. (B) Quantification of relative expression of tuberin (across 3 technical replicates using these lines). (C) Representative immunofluorescence images of induced pluripotent stem cells showing expression of TUBA (magenta) and Hoechst (teal) to identify DNA to identify mitotic cells. Scale bars = 50 µm (D) Quantification showing average percentage of mitotic cells that have the multipolar division by genotype. N = 3 independent replicates per genotype. No significance [paired students t test]. Error bars = SD.

### Patient-derived mutant TSC2 exhibits a dominant negative effect

Since the multipolar division phenotype was observed in the iPSCs with the mutant tuberin but not the *TSC2* KO iPSCs, we next tested whether the mutant version of TSC2 that is expressed in patient-derived iPSCs exerts a dominant negative effect. We performed a transient siRNA knock down in the patient mutant lines to remove the mutant TSC2 protein. Loss of TSC2 protein was observed in all iPSCs treated with siRNA against *TSC2* (Figure 3A). The proportion of multipolar division significantly decreased between the scramble control and siTSC2 treated patient TSC2 +/LOF iPSCs (p=0.0307) (Figure 3C). The percentage of cells in mitosis significantly increased in the *TSC2 +/LOF* siTSC2 treated iPSC compared to scramble control cells (p=0.0095) (Figure 3B). Interestingly, the average rate of mitosis and proportion of multipolar divisions did not significantly differ between scramble control and siTSC2-treated cells in the patient wild type. The same is true in scramble control and siTSC2-treated *TSC2* LOF/LOF iPSCs (Figure 3B, C).

**Figure 3:**
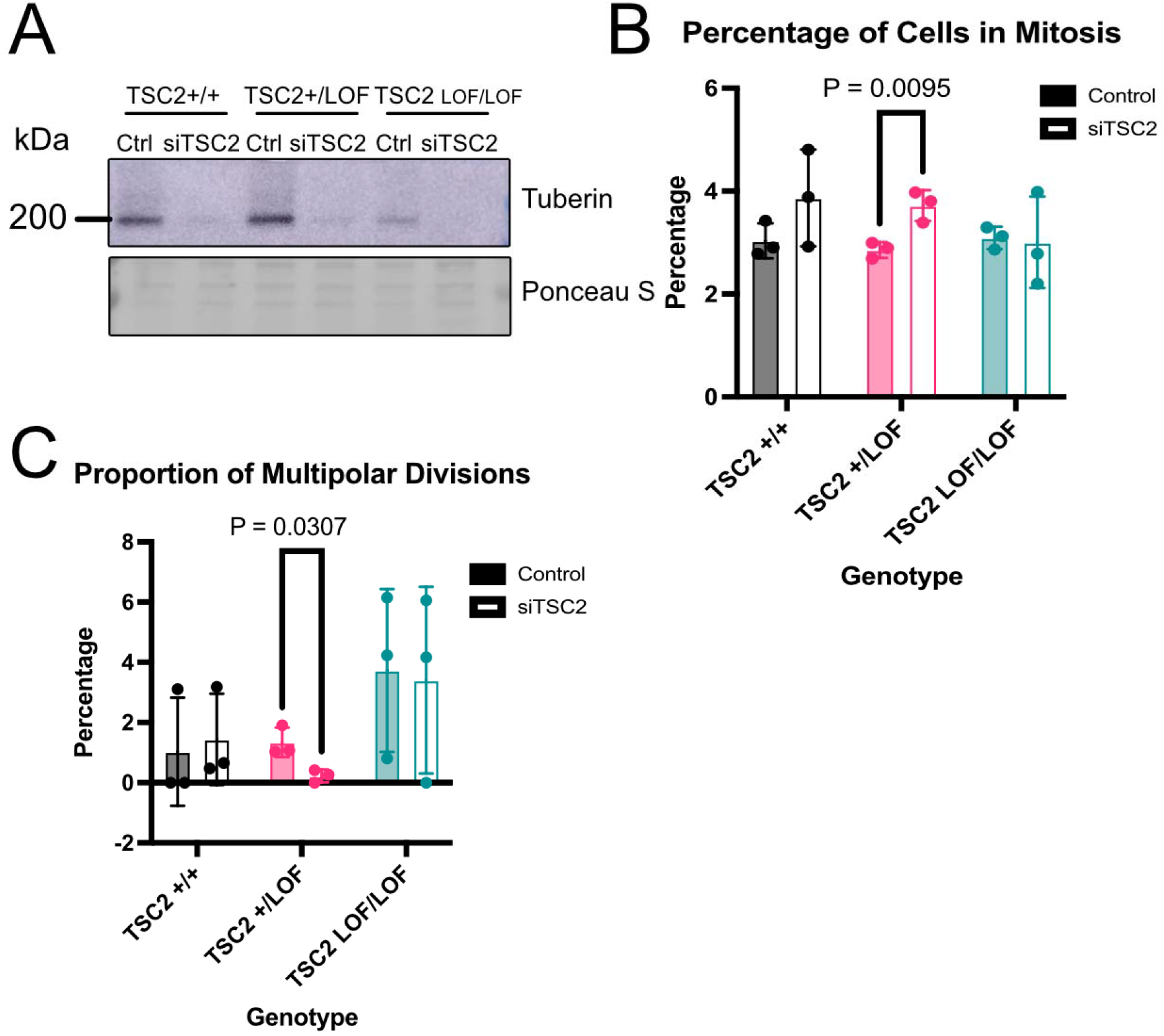
siTSC2 induced pluripotent stem cells have multipolar mitotic cells. (A) Representative immunoblot of induced pluripotent stem cells showing expression loss of Tuberin in siTSC2 iPSCs. (B) Quantification showing average percentage of cells in mitosis per genotype and treatment. N = 3 independent replicates per genotype and treatment, p = 0.0095 [paired students t test]. Error bars = SD. (C) Quantification showing average percentage of mitotic cells that have the multipolar division per genotype and treatment. N = 3 independent replicates per genotype and treatment, p = 0.0307 [paired students t test]. Error bars = SD.

## DISCUSSION

TSC is a prototypical neurogenetic disorder whose study has led to many insights to the normal process of brain development as well as the pathogenesis of epilepsy, autism, and intellectual disabilities. While the disorder is caused by mutations in *TSC1* or *TSC2* genes, patients with *TSC2* mutations typically have more severe symptoms (Ess, 2010; Sancak *et al*., 2005). In this study, we chose to use an isogenic allelic series of iPSCs from a patient with a *TSC2* mutation (Sundberg *et al*., 2018). This particular *TSC2* mutation (c.5238_5255del, p. (His1746_Arg1751del of 6 amino acids in exon 41) has been frequently reported. In fact, in multiple studies profiling mutations in patients with TSC throughout the world, this same 6 amino acids deletion was found to be the most common mutation (Dabora *et al*., 2001; Jones et al., 1999; Reyna-Fabián et al., 2020; Rosengren et al., 2020; Wang et al., 2013). The impact from this specific mutant microdeletion allele thus has important consequences for patients with TSC globally.

Approximately 90% of TSC patients experience epilepsy which is thought to originate from cortical tubers (Thiele, 2004). A diagnostic characteristic of cortical tubers is the presence of giant cells. Prior studies of these lesions include reports that a subpopulation are multinucleated (Crino, 2013; Feliciano, 2020; Mizuguchi, 2007; Zarei *et al*., 2002). Multinucleated cells can arise from several different mechanisms. One such mechanism is multipolar division, in which a cell has more than two mitotic spindle poles (THERMAN and TIMONEN, 1950). A cell undergoing multipolar division may then not complete cytokinesis, resulting in a single cell with more than one nucleus (Bruce Alberts, 2008). As multinucleated giant cells are pathognomonic for cortical tubers, an unanswered question is whether such early cell division abnormalities during early brain development underlie the formation of cortical tubers in patients with TSC. We found that iPSC cultures with both heterozygous and homozygous *TSC2* mutation include a subpopulation of cells undergoing multipolar division, indicating a role for tuberin in proper division during mitosis. A possible contributing mechanism to the formation of multinucleated giant cells in cortical tubers from mutant tuberin may thus be abnormal mitosis with failed cytokinesis through multipolar division. Further study will be required to determine if such a mechanism contributes to the pathogenesis of TSC.

When we investigated *TSC2* KO iPSCs, which lack all tuberin protein, we did not find any nuclei undergoing multipolar division. As the complete absence of tuberin did not result in multipolar cells, expression of a mutant tuberin protein lacking 6 amino acids seems to be required to trigger multipolar division. To test this possibility, we knocked down tuberin with siRNA and saw that in the TSC2 heterozygous iPSCs, upon treatment of siTSC2, there was an increase in the percentage of cells in mitosis and a decreased incidence of multipolar division. These findings indicate a dominant negative effect of the mutant tuberin against the wild type tuberin. The patient wild type and *TSC2* complete *LOF/LOF* iPSCs did not show the dominant negative effect likely due to neither genotype having both a wild type copy and a mutant copy of tuberin. Dominant negative activity from a heterozygous TSC2 mutation, as indicated by these data, could indicate a novel mechanism of disease that has not been previously widely considered. Future studies can determine if any other TSC2 mutations may also lead to an increase in multipolar dividing cells.

The multipolar phenotype is likely mTORC1-independent indicated by rapamycin treatment not effecting the rate of multipolar division in each genotype. Previous studies on tuberin function have primarily focused on mTORC1-dependent phenotypes, but more investigation into mTORC1-independent tuberin functions is strongly needed as some aspects of the TSC phenotype appear to be resistant to treatment with rapamycin or structurally related drugs (Bissler et al., 2008; French et al., 2016; Krueger et al., 2013; McCormack et al., 2011; Valianou et al., 2019).

The TSC heterodimer is composed of hamartin and tuberin (Tee *et al*., 2003; van Slegtenhorst *et al*., 1997). Hamartin has been shown to localize to the centrosome. Hamartin and the entire TSC heterodimer was found to interact with PLK1 (polo-like kinase 1), a kinase involved in the duplication of centrioles (Astrinidis et al., 2006; Shukla et al., 2015). Additionally, TACC3, a kinase involved in centrosome activity and formation of microtubules during cell division, has been shown to phosphorylate tuberin localizing tuberin to the centrosomes (Gergely et al., 2000; Gómez-Baldó et al., 2010; Peset and Vernos, 2008). These data, combined with the findings from this study, indicate a role for tuberin and the TSC complex in centriole duplication, suggesting a possible mechanism that may underlie multipolar division. We anticipate that the results presented here will be a catalyst for further analysis of additional TSC patient-derived iPSCs for the presence of multipolar cells and the mechanisms behind their origin.

## EXPERIMENTAL PROCEDURES

### Resource Availability

#### Corresponding Author

Further information and requests for resources and reagents should be directed to and will be fulfilled by the co-corresponding author, Kevin Ess.

#### Materials availability

This study did not generate new unique reagents.

### hiPSC Cell Culture

iPSCs were grown as colonies on Matrigel (Corning) coated 6 well plates or glass bottom 35mm plates (Cellvis) in mTeSR1 medium (StemCell Tech), replaced daily, maintained at 37°C and 5% CO_2,_ and passaged as needed with ReLSR (StemCell Tech).

### Whole Cell Extract

Cells were washed with 1X PBS with 10uM sodium vanadate. Cells were lysed using lysis buffer (1% Triton X-100 in STE [100mM NaCl, 1mM EDTA, 10mM Tris pH 8.0], PhosSTOP (Sigma Aldrich), PIC (Sigma Aldrich), 50uM MG132, 1uM PMSF) added directly to the plate followed by scraping of the cells. Lysate was then sonicated for 3 seconds on power 3 at 4°C. Lysate was centrifuged for 15 minutes at 16,000 x g at 4°C. Supernatant was kept at -80°C until further analysis.

### Immunoblotting

Samples were prepared by mixing whole cell extracts with 4X SDS (find company) and then boiled for 5 minutes at 105°C. Gel electrophoresis was performed on a 4-12% Bis-Tris gel (ThermoFisher) with 1X NuPage MOPS running buffer (ThermoFisher) at a constant voltage of 140V for 90 minutes. Proteins were transferred to a PVDF membrane in transfer buffer on ice in the cold room (4°C) overnight at a constant current of 33 mA. Post-transfer, the membrane was stained with Ponceau S solution (Sigma Aldrich) for 5 minutes to determine total protein loaded into gel lanes.Membranes were washed 3X in ddH2O. Membranes were blocked in 5% non-fat milk in Tris-buffered saline with 0.1% Tween 20 (TBS-t) for 1 hour at room temperature with agitation. All primary and secondary antibodies were diluted in TBS-t with 5% non-fat milk, and the membrane was washed 3X with 0.1% TBS-t afterwards. Antibodies are listed in the table 1. Primary antibodies were incubated overnight at 4°C with agitation. Secondary antibodies were incubated for 1 hour at room temperature with agitation. The membrane was imaged using ECL developing reagents (ThermoScientific) and CCD Imager – AI600 (General Electric).

**Table 1:** Antibodies used in these experiments.

| Antibody | Manufacturer | Catalog | Dilution | Application |
| --- | --- | --- | --- | --- |
| Alpha Tubulin<br>(TUBA) | Invitrogen | MA1-80017 | 1:500 | Immunostaining |
| GAPDH | Cell Signaling<br>Technologies | 2118 | 1:2000 | Western Blot |
| TUBERIN<br>(TSC2) | Cell Signaling<br>Technologies | 4308 | 1:2000 | Western Blot |
| Goat anti-Rat<br>IgG, Alexa<br>Fluor 594 | Invitrogen | A-11007 | 1:1000 | Immunostaining |
| Goat anti-Rat<br>IgG, Alexa<br>Fluor 568 | Invitrogen | A-11077 | 1:1000 | Immunostaining |
| Goat anti-rabbit IgG, HRP | Invitrogen | 65-6120 | 1:5000 | Western Blot |

### Immunofluorescence

All cells were fixed by incubation in 100% methanol for 10 minutes at -20°C. Fixed samples were blocked with blocking buffer [PBS, 1% Normal Donkey Serum, 1% BSA, 0.1% Triton X-100] for 1 hour at room temperature. Primary antibodies were diluted in blocking buffer and then incubated overnight at 4C. Secondary antibodies were diluted in blocking buffer and then incubated 1 hour at room temperature in the dark. Antibodies are listed in the table 1. Images were acquired using a Prime 95B camera mounted on a Nikon spinning disk microscope using a Plan Apo Lambda 20x objective lens. The software used for image acquisition and reconstruction were NIS-Elements Viewer (Nikon) and ImageJ (FIJI).

### Image Analysis

Images were blinded prior to scoring. CellProfiler was then used to quantify the total number of cells in each image. Subsequently, cells were manually identified to be in prophase, metaphase, anaphase, or telophase. The number of cells with more than two mitotic spindles was also recorded. Two-way ANOVA and student t-tests were run using Prism (GraphPad) on the collected data.

### siRNA transfection of human iPSCs

Cells were grown as described previously. When iPSCs reach 50% confluency, 1uM siRNA (Horizon Accell SMARTpool) in DMEM/F12 (Gibco) was added to the cells. siRNA incubated on the cells for 24 hours. Media was changed to mTeSR1 after 24 hours and cells were allowed to grow normally for 48 more hours. iPSCs were then fixed or lysed.

## Acknowledgements

We would like to thank Asa Brockman (Ihrie lab) for the introduction and troubleshooting of CellProfiler workflows, as well as Ihrie, Ess, and Irish lab members for thoughtful discussions of data. We would also like to thank the Vanderbilt Cell Imaging Shared Resource and Nikon Center of Excellence. Experiments/Data analysis/presentation were performed in part through the use of the Vanderbilt Cell Imaging Shared Resource (supported by NIH grants CA68485, DK20593, DK58404, DK59637 and EY08126). This research was supported by awards R01NS118580 (RAI, KCE), R01NS118580-S1 (MBLC), and T32 HD007502 (MBLC).

## Contributions

Conceptualization, Funding Acquisition – MBLC, RAI, KCE. Formal Analysis, Investigation, Software – MBLC, RBM. Methodology – MBLC, MS, KCE. Resources – MS, MuS, LA, RAI, KCE. Validation, Visualization – MBLC. Writing – Original Draft – MBLC, RBM. Writing – Review and Editing – MBLC, MuS, LA, RAI, KCE

